# *TaPGS1* Driven Flavonol Accumulation Delays Endosperm Cellularization and Enlarges Wheat Grains

**DOI:** 10.1101/2025.06.17.660139

**Authors:** Xiaojiang Guo, Xin Liu, Yumeng Jin, Shuyu Zhao, Wangzhuang Liang, Maolian Li, Mengping Cheng, Huixue Dong, Qian Chen, Zhongxu Chen, Jirui Wang

## Abstract

Flavonols play a crucial role in seed development by regulating multiple physiological processes, including seed coat pigmentation, dormancy, fertilization, and endosperm formation. Notably, flavonols influence the polar transport of auxin, thereby affecting seed growth dynamics. In our previous work, we found that overexpression of the wheat bHLH transcription factor TaPGS1 results in increased grain size; however, the underlying mechanism remained unclear. In this study, metabolomic and transcriptomic analyses of *TaPGS1* overexpressing wheat lines revealed enhanced flavonol accumulation and upregulation of key flavonol biosynthetic genes. Further investigations suggested that flavonol accumulation in the seed coat may disrupt auxin transport, leading to localized auxin buildup, delayed endosperm cellularization, and an increase in endosperm cell number. These changes collectively contribute to grain enlargement. Our findings uncover a *TaPGS1* flavonol regulatory module that links auxin distribution to endosperm development and seed size control in wheat.

## 1. Introduction

Seed size is a crucial agronomic trait that determines crop yield potential, controlled by the growth and development of the seed coat, embryo, and endosperm (Xu and Zhang 2023). The endosperm is the primary nutrient storage organ in wheat, constituting over 80% of the seed’s dry weight in a fully developed mature wheat seed (Egli et al. 2017). and is therefore the major sink for starch and storage proteins (Liu et al. 2022). The degree of endosperm development determines the size and weight of wheat grains (Ji et al. 2022), directly affecting the yield and quality of wheat grains. Endosperm development in cereals follows the nuclear type of endosperm development, where initial endosperm nuclei divide without immediate cytokinesis, forming a coenocyte (early endosperm) (Olsen 2004). As development progresses, the endosperm enters the cellularization stage, where the endosperm nuclei are partitioned by newly formed cell walls to create cellularized endosperm cells (Costa et al. 2004; Olsen 2001). Cellularization largely fixes the internal volume of the grain, so its timing strongly influences final grain size (Berger et al. 2006; Bin Zhang et al. 2020; Zhou et al. 2009).

Auxin acts as a developmental timer for that governs the switch from the coenocytic (syncytial) endosperm to the cellularized endosperm stage. High auxin concentrations in the coenocytic endosperm are sustained by polar export to the surrounding seed coat; when auxin falls below a threshold, cellularization is triggered (Figueiredo et al. 2016; Batista et al. 2019). Flavonoids have been reported to regulate auxin transport or local concentration, affecting the timing of endosperm cellularization (Besseau et al. 2007; Scott et al. 2013). For instance, kaempferol and quercetin, modulate auxin efflux by inhibiting PIN- and P-glycoprotein (PGP) transporters and their kinase regulator PINOID (PID) (Besseau et al. 2007; Scott et al. 2013; Blakeslee et al. 2007; Teale et al. 2020). Genetic studies in *Arabidopsis* corroborate this link: the *tt4* mutant (defective in Chalcone synthase, CHS) displays elevated polar auxin transport, whereas overexpression of *TT3* (DIHYDROFLAVONOL 4-REDUCTASE, *DFR*) or *TT7* (FLAVONOID 3□-HYDROXYLASE, *F3*□*H*) reduces it (Buer and Muday 2004; Peer et al. 2004; Yin et al. 2013). Because both flavonol and auxin pathways branch from phenylalanine, feedback between their metabolic nodes may fine-tune auxin distribution (Sébastien et al. 2007; Dixon and Paiva 1995; Orlova et al. 2006). Yet it remains unclear how this metabolic–hormonal interaction governs the timing of endosperm cellularization and, by extension, grain size in wheat.

We previously characterized *TaPGS1* (*Triticum aestivum Positive Regulator of Grain Size 1*), a bHLH transcription factor homologous to *Arabidopsis TT8* (*TRANSPARENT TESTA 8*) and rice *OsRc* (*Oryza sativa, red pericarp gene*). Whereas *TT8* and *OsRc* influence seed-coat pigmentation and dormancy (Gu et al. 2011; Nesi et al. 2000). *TaPGS1* overexpression enlarges wheat grains without altering pericarp color and is accompanied by increased expression of flavonol-biosynthetic genes(Guo et al. 2022). These observations suggest that *TaPGS1* may remodel flavonol metabolism to regulate auxin dynamics and cellularization, but the underlying mechanism is unknown.

Here, we integrate transcriptomic and metabolomic profiling of *TaPGS1* overexpression lines to identify differentially expressed genes and metabolites in the flavonol pathway. We visualize flavonol and auxin localization in developing seeds, assess the effect of exogenous kaempferol on cellularization, and test direct TaPGS1 targets using dual-luciferase, yeast two-hybrid, and BiFC assays. Our study uncovers a TaPGS1-centred regulatory module that links flavonol accumulation to auxin-mediated control of endosperm cellularization, providing mechanistic insight into the genetic improvement of wheat grain size.

## 2. Materials and methods

### 2.1 Plant materials and phenotype

We used common wheat (*Triticum aestivum*) cultivar Fielder and two *TaPGS1* overexpression lines (OE-166-6 and OE-166-39) in the Fielder background as study materials. Wild-type Fielder and the overexpression lines were grown at Wenjiang, China (30°72′ N, 103°87′ E). Field traits, plant height, spike length, tiller number, effective spike number, and grain number per spike, were recorded. Spikes at the soft dough stage were harvested, air-dried, and manually threshed, and grain moisture was adjusted to ∼10 % with a near-infrared analyzer (DA 7250, PerkinElmer). For germination tests, fifty seeds per line (three replicates) were sown on moist filter paper, germination was scored daily for seven days, and germination percentage was calculated. Mature seeds from the overexpression lines were harvested, naturally dried to ∼10 % moisture, and evaluated for seed length, width, and color. (five replicates per line). Other wheat plants were grown in a greenhouse under a 16 h light / 8 h dark photoperiod at 20 °C and 55–60 % relative humidity, while *Nicotiana benthamiana* plants were cultivated in a growth chamber under the same photoperiod at 25 °C.

### 2.2 RNA-seq differential expression analysis and quantitative real-time PCR

Total RNA was extracted from developing seeds of *TaPGS1* overexpression lines and wild-type Fielder grown in a greenhouse at 5, 10, 15, and 20 days post-anthesis (DPA). Strand-specific cDNA libraries were constructed using the standard Illumina protocol and sequenced on an Illumina HiSeq 2500 platform. Raw reads were quality-trimmed using the FASTX toolkit and mapped to the wheat reference genome (Chinese Spring, IWGSC RefSeq v1.0) with TopHat v2.0. Transcript assembly and abundance quantification were performed using Kallisto, and differentially expressed genes were identified. Heat maps and preliminary functional annotations were generated using GraphPad Prism v8.1.

For quantitative real-time PCR (qPCR), total RNA was reverse-transcribed using the PrimeScript^™^ 1st Strand cDNA Synthesis Kit (Takara, Beijing, China). qPCR reactions were conducted withSYBR Premix Ex Taq II (Takara, Beijing, China) on a CFX96 Touch^™^ Real-Time PCR Detection System (Bio□Rad) under the following thermal profile: initial denaturation at 95□°C for 1□min, followed by 40 cycles of 95□°C for 5□s and 60□°C for 30□s. Relative transcript levels were normalized to internal reference genes as described by Long (Long et al. 2010). Primer sequences are listed in Table□ S1.

### 2.3 Metabolomic analysis

Metabolomic profiling was carried out on 5-DPA seeds of *TaPGS1* overexpression lines and wild-type Fielder. Crude extracts were analyzed by LC–MS, and raw data were processed in Compound Discoverer (CD) version 3.3 (Thermo Fisher Scientific) for peak detection, retention-time alignment, and feature deconvolution. Molecular features were matched against the mzCloud, mzVault, and Mass List databases with a mass-accuracy window of ≤5 ppm to obtain putative metabolite identifications. Peak areas integrated in CD 3.3 were normalized to the total ion current (TIC) of each sample and used to represent the relative abundance of metabolites across samples. Differential metabolites were defined by |log□ fold change| ≥ 1 and FDR < 0.05 and visualized as heat maps in GraphPad Prism V8.1. Finally, changes in flavonol–auxin pathway metabolites were compared with differentially expressed genes identified from the transcriptome to explore coordinated regulation at the metabolic and transcriptional levels.

### 2.4 Flavonol fluorescence staining localization

Fresh seeds at 5 DPA were hand-sectioned and incubated for 5 min at room temperature in diphenylboric acid 2-aminoethyl ester (DPBA) staining solution. After staining, the sections were rinsed twice with distilled water, mounted on microscope slides, and examined immediately with a fluorescence microscope (BX61, Olympus). Under DPBA staining, kaempferol fluoresces green and quercetin fluoresces yellow, allowing visualization of flavonol accumulation in the endosperm (Nguyen, et al. 2020).

### 2.5 Auxin localization (immunohistochemistry)

Seeds of wild-type Fielder and *TaPGS1* overexpression lines at 5 DPA were used for immunolocalization of indole-3-acetic acid (IAA) following a whole-mount immunohistochemical method. Samples were vacuum-infiltrated with EDC/NHS buffer containing 0.1% Tween-20, then fixed overnight at 4□°C in 4% PFA with 10% DMSO and 3% NP-40. After dehydration through a graded methanol–ethanol series, tissues were cleared at 65□°C and permeabilized with 100 μg/mL proteinase K. Membrane permeability was further enhanced with 10% DMSO, and non-specific binding was blocked with 3% BSA. Samples were then incubated with an anti-IAA primary antibody (1:100) followed by an HRP-conjugated secondary antibody (1:500). Signal was visualized with 3,3 ′-diaminobenzidine (DAB), enhanced by methanol dehydration, and tissues were cleared in chloral hydrate for observation under a bright-field optical microscope (Wang, et al. 2022).

### 2.6 Semi-thin sectioning

Seeds from greenhouse-grown wild-type Fielder and *TaPGS1* overexpression lines were harvested from 0 to 30 days after flowering and immediately fixed under vacuum in aldehyde-based fixative for 12–24 h. After three rinses in phosphate buffer (10–20 min each), samples were dehydrated through an ethanol series of 25 %, 40 %, 60 %, 70 %, 80 %, 90 %, and 100 % (30 min at each concentration). Dehydrated tissues were then infiltrated with epoxy resin solutions of 25 %, 50 %, 75 %, and 100 %, allowing at least 2 h for each step and holding the final 100 % resin overnight (≈12 h). Specimens were embedded in fresh epoxy resin and polymerized at 65 °C for 12–24 h. Semi-thin sections (EM UC7, Leica) were cut on an ultramicrotome, stained with 0.1 % toluidine blue, and examined under a microscope (Axio lmager M2, Zeiss) to document endosperm cellularization and cell-division patterns. For seeds collected 4–7 days after flowering, the cellularization status was scored and the proportion of fully cellularized seeds was calculated for each developmental stage.

### 2.7 Luciferase (LUC) reporter assay

Promoter fragments of *TaDFR* and *TaF3H* were amplified from wheat genomic DNA and inserted into pGreenII-0800-LUC, generating the reporter constructs *TaF3H*pro:LUC and *TaDFR*pro:LUC. The *TaPGS1* coding sequence was cloned into pGreenII-62SK to obtain the effector construct 35Spro:TaPGS1 (primer sequences are listed in Table S1). *Agrobacterium tumefaciens* GV3101 carrying 35Spro:TaPGS1 was co-infiltrated with either *TaF3H*pro:LUC or *TaDFR*pro:LUC into fully expanded leaves of three-week-old *Nicotiana benthamiana*; leaves co-infiltrated with the empty pGreenII-62SK vector and the same reporters served as negative controls. At 36–48 h post-infiltration, firefly (LUC) and Renilla (REN) luciferase activities were measured with a Dual-Luciferase Reporter Assay kit (Promega, Madison, USA), luminescence images were captured with Biorad ChemiDoc MP Imaging System (Bio-Rad), and relative promoter activity was expressed as the LUC/REN ratio. Each treatment was assayed with three biological replicates (Hellens et al. 2005).

### 2.8 Yeast two-hybrid assay and bimolecular fluorescence complementation

Full-length coding sequences of *TaPGS1*, *TaMYB*, and *TaWD40* (gene IDs in Table S4) were amplified from Fielder cDNA. *TaPGS1* was inserted into the pGADT7 activation-domain (AD) vector, whereas *TaMYB* and *TaWD40* were cloned into the pGBKT7 DNA-binding-domain (BD) vector. Each BD construct (BD-MYB or BD-WD40) was co-transformed with AD-TaPGS1 into Saccharomyces cerevisiae strain AH109, and empty AD and BD vectors served as negative controls. Transformants were selected on SD/-Leu/-Trp medium and then transferred to SD/-Leu/-Trp/-His medium to test interaction-dependent growth; colonies were scored after 36–48 h.

For bimolecular fluorescence complementation (BiFC), the *TaPGS1* coding sequence was fused to the N-terminal fragment of YFP in vector 35S-SPYNE, and full-length TaMYB or TaWD40 was fused to the C-terminal fragment of YFP in vector 35S-SPYCE. Sequenced constructs were introduced into *Agrobacterium tumefaciens* GV3101. Cultures containing TaPGS1-YNE were mixed 1: 1 (v/v) with either TaMYB-YCE or TaWD40-YCE and infiltrated into the abaxial side of fully expanded leaves of three-week-old *Nicotiana benthamiana* using a needle-less syringe. After 36–48 h, YFP fluorescence was observed with a laser-scanning confocal microscope using 514 nm excitation to confirm protein–protein interactions in planta.

### 2.9 Flavonol (kaempferol) treatment

Seeds of wild-type Fielder were collected 4 days after flowering and placed on half-strength MS medium supplemented with 0, 10, 25, or 50 µM kaempferol. After 24 h and 48 h of incubation, seeds from each treatment (three biological replicates) were fixed in paraformaldehyde solution and processed for semi-thin sectioning as described in Section 2.6.

## 3. Results

### 3.1 *TaPGS1* overexpression enlarges wheat grains without major agronomic trade-offs

The homologous genes of *TaPGS1* in *Arabidopsis* (*TT8*) and rice (*OsRc*) both affect grain color, and *OsRc* also regulates seed dormancy and weight in rice(Gu et al. 2011; Nesi et al. 2000). In wheat, overexpression of *TaPGS1* results in increased seed size. Compared to the Fielder variety, the *TaPGS1* overexpression lines exhibit significantly higher thousand-grain weight, grain length, and grain width phenotypes in field conditions (Fig. 1.A, D-F). Further analysis of anthocyanin accumulation in grains treated with NaOH from red-skinned cultivar Chuanmai 104, wild-type Fielder, and *TaPGS1* overexpression lines (OE-166-6 and OE-166-39) in the Fielder background revealed no color difference between the *TaPGS1* overexpression grains and the wild-type Fielder grains. Only the red-skinned Chuanmai 104 showed higher grain permeability and deeper color (Fig.1.B). Subsequently, germination vigor of seven-day-old seeds from wild-type Fielder and *TaPGS1* overexpression lines was assessed, showing no significant difference in germination rates between the wild-type and overexpression lines (Fig. 1.C). Additionally, field phenotypic traits, including plant height, spike length, tiller number, effective spike number, and grains per spike, were evaluated. Results indicated that, except for the OE-166-39 line, which exhibited increased plant height and tiller number compared to Fielder, there were no significant differences in other traits (Fig. S1.A-E). These findings suggest that overexpression of *TaPGS1* primarily affects seed size with minimal impact on other agronomic traits in wheat.

**Fig. 1.**
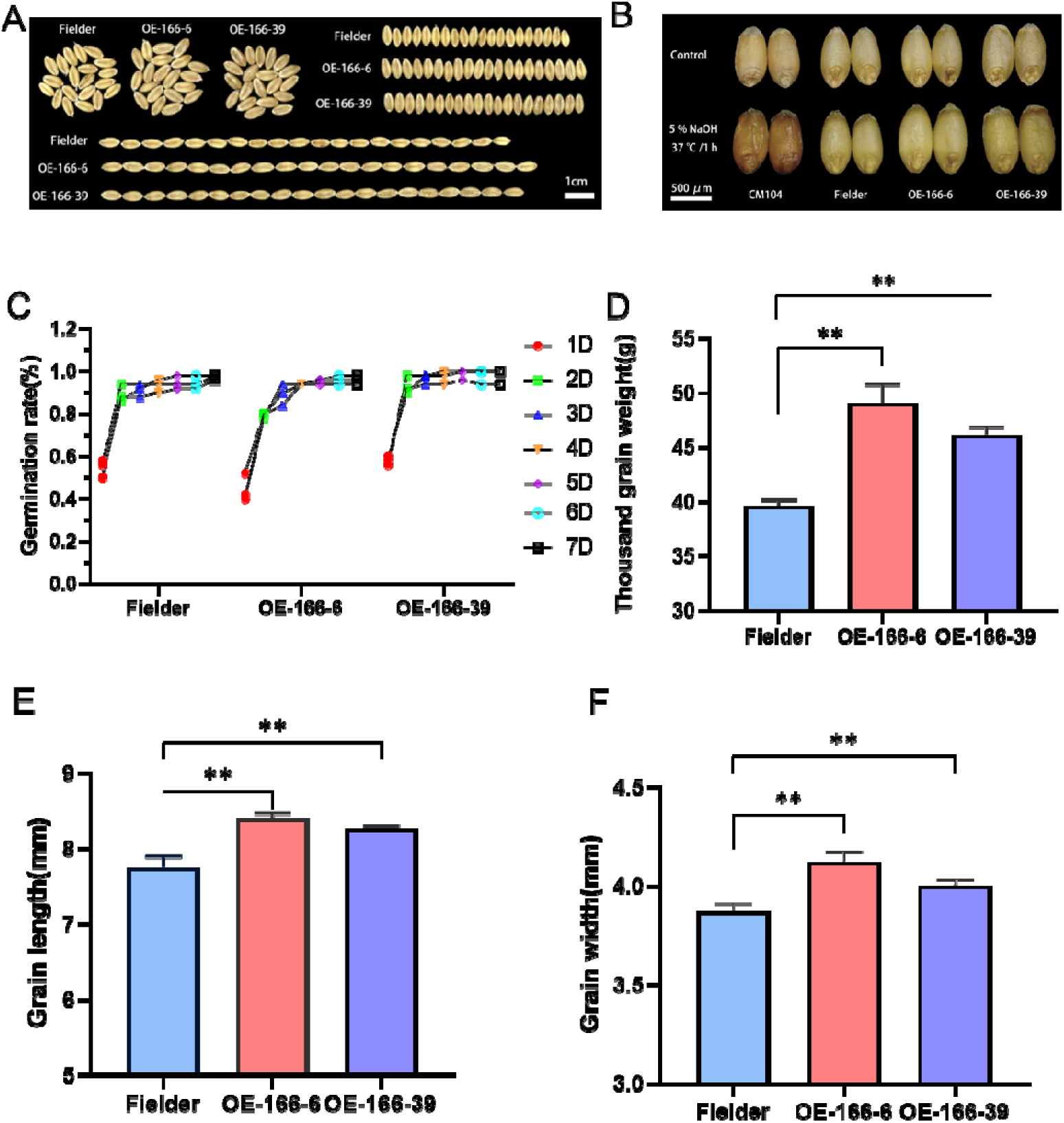
Analysis of wheat grain phenotype and key agronomic traits in the field. **A**. Comparison of grain size between wild-type Fielder and *TaPGS1* overexpression wheat. **B**. Comparison of grain permeability between wild-type Fielder, *TaPGS1* overexpression lines, and red-skinned cultivar Chuanmai 104 (CM104) after NaOH treatment. **C**. Comparative analysis of seven-day germination vigor between wild-type Fielder and *TaPGS1* overexpression lines. D-F. Field analysis of thousand-grain weight (**D**), grain length (**E**), and grain width (**F**) between wild-type Fielder and *TaPGS*1 overexpression lines (* *P* < 0.05; ** *P* < 0.01).

### 3.2 *TaPGS1* overexpression enhances flavonol-pathway gene expression and metabolite levels

The total metabolites in seeds of wild-type Fielder and *TaPGS1* overexpression wheat five days post-anthesis were measured. Metabolomic analysis revealed that flavonol pathway metabolites were the most significantly detected (Fig. 2A). Further analysis of these 42 enriched metabolites showed that the levels of naringenin, naringin, and kaempferol were higher in *TaPGS1* overexpression lines compared to the wild type (Fig. 2B, Table S2). Concurrently, through transcriptome sequencing, this study obtained the expression levels (Transcripts Per Million, TPM values) of whole genome genes in seeds from wild-type Fielder and *TaPGS1* overexpression lines at 5, 10, 15, and 20 days post-anthesis (Table S3). Differential expression analysis and qPCR revealed a significant upregulation of several key enzyme genes in the flavonoid biosynthesis pathway, including *F3H*, *DFR*, *FLS* (*Flavonol Synthase*) and *ANS* (*Anthocyanidin Synthase*), in the *TaPGS1* overexpression lines (Fig. 2C; Fig. S2). These findings are consistent with the observed upregulation of flavonol pathway metabolites in the metabolomic analysis (Fig. 2A, B, C; Fig. S2).

**Fig. 2.**
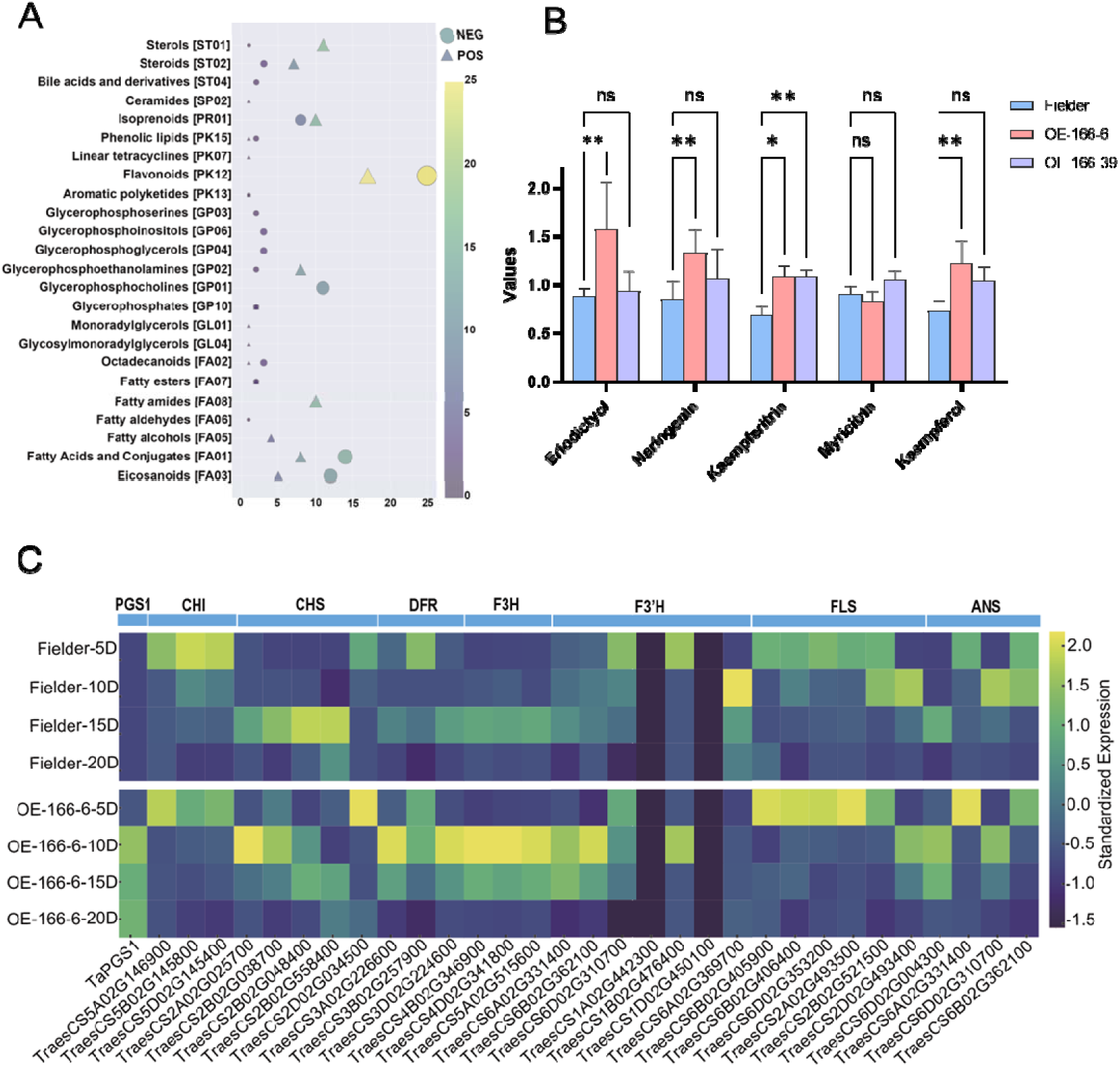
Metabolomic and transcriptomic analysis of Fielder and *TaPGS1* overexpression lines. **A**. Differential metabolite enrichment analysis. **B**. Analysis of key metabolites in the flavonol biosynthesis pathway. Values represent the normalized peak area (arbitrary units) from five biological replicates. These values reflect the relative metabolite levels after total ion current normalization. (mean ± SD; * *P* < 0.05; ** *P* < 0.01). **C**. Transcriptomic analysis of the expression of genes related to flavonoid biosynthesis in wild-type Fielder and *TaPGS1* overexpression lines.

Based on these findings, we performed section and staining analysis on seeds from wild-type Fielder and *TaPGS1* overexpression lines 5 DPA. Seeds were sectioned and stained with DPBA, followed by observation under a fluorescence microscope. The results showed that the accumulation of the flavonol metabolite kaempferol in *TaPGS1* overexpression seeds was significantly higher than in Fielder seeds, with fluorescence intensity in OE-166-6 and OE-166-39 being 2.0 and 1.3 times higher, respectively, than in Fielder. In contrast, the levels of another metabolite, quercetin, did not significantly differ between *TaPGS1* overexpression lines and Fielder (Fig. 3A, B). These results indicate that overexpression of *TaPGS1* leads to increased expression of flavonol pathway genes and accumulation of the flavonol metabolite kaempferol in wheat seeds.

**Fig. 3.**
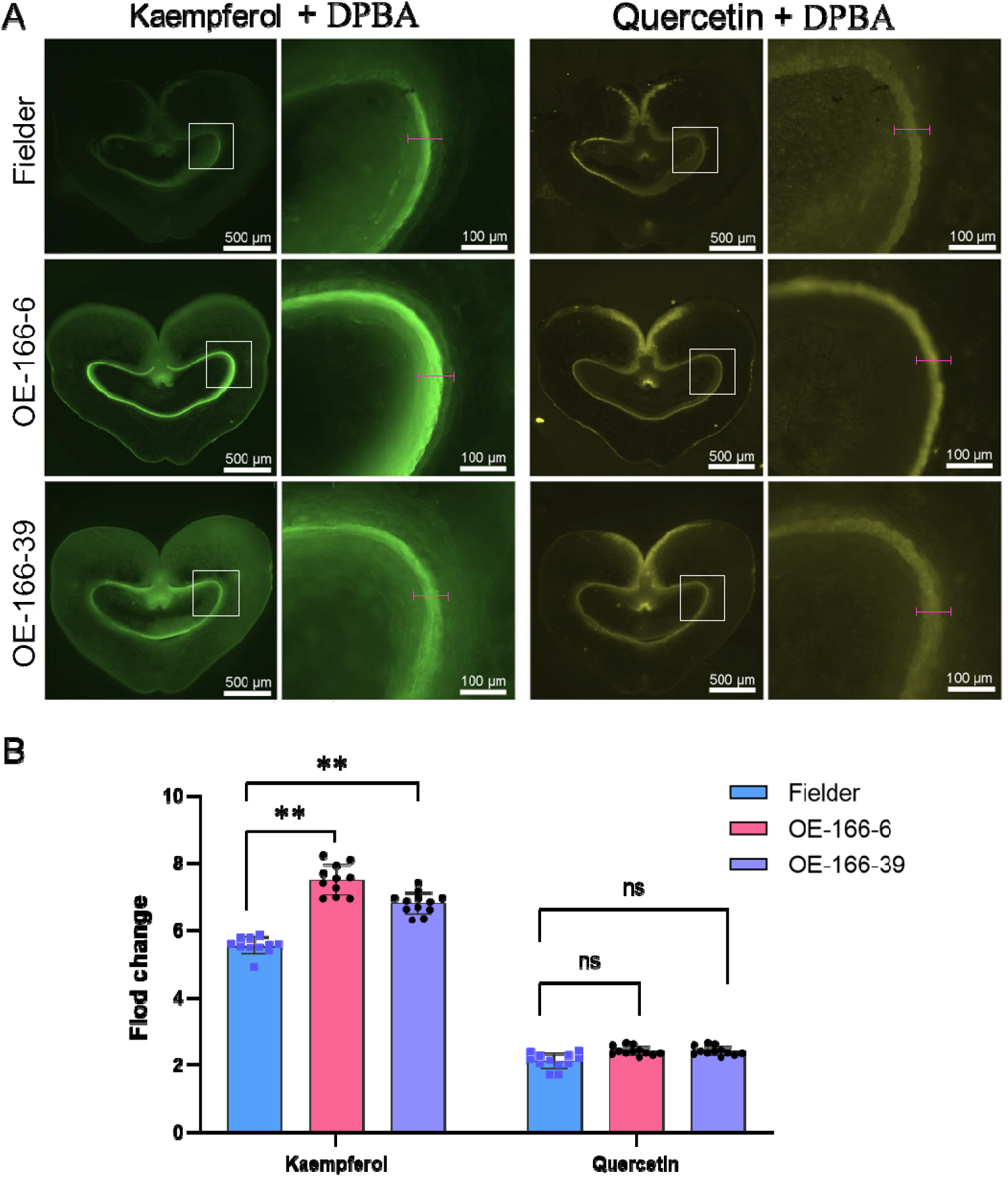
Localization and accumulation of flavonols in wheat seeds. **A**. Fluorescence observation of seeds five days after flowering stained with DPBA (green fluorescence indicates kaempferol, yellow fluorescence indicates quercetin). **B**. Quantitative analysis of fluorescence intensities of kaempferol and quercetin from A (n = 10; * *P* < 0.05; ** *P* < 0.01). The white box indicates the enlarged display area. The red line segment represents the data collection area.

### 3.3 *TaPGS1* overexpression elevates auxin and delays endosperm cellularization

The accumulation of flavonols affects auxin transport; in flowering plants, auxin is transported from the endosperm to the seed coat post-fertilization. If this transport is impaired, endosperm cellularization is affected (Besseau et al. 2007; Scott et al. 2013; Figueiredo et al. 2016). To verify auxin accumulation in the inner seed coat, we employed an in situ localization method to observe auxin distribution in 5-day-old wheat seeds. The results indicated that auxin accumulates in the inner seed coat in the *TaPGS1* overexpression lines (Fig.4A, B).

**Fig. 4.**
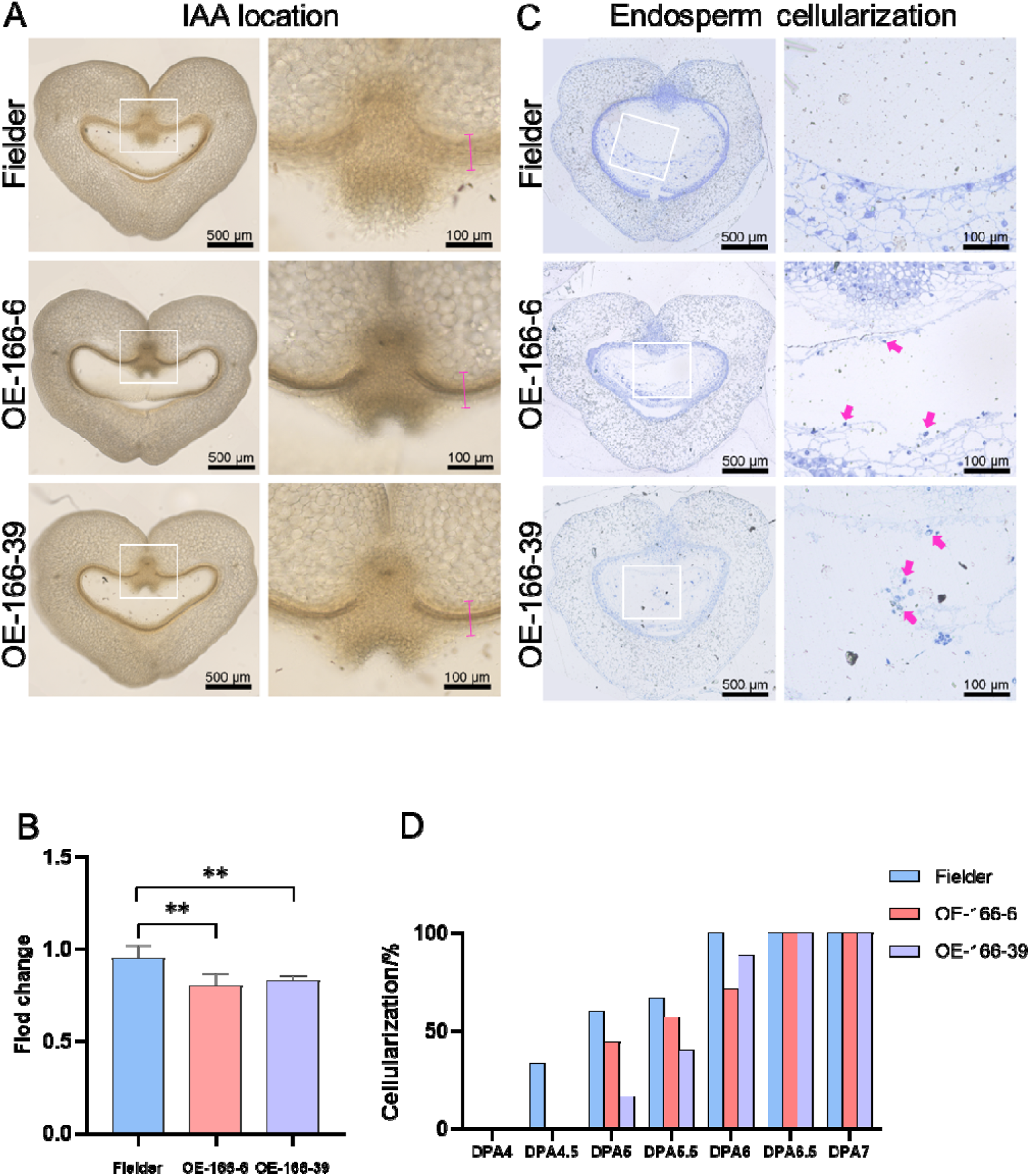
Analysis of auxin localization and endosperm cellularization phenotype in TaPGS1 overexpression lines. **A.** Auxin accumulation in the seeds of wild-type Fielder and *TaPGS1* overexpression lines at 5 DPA. The white box indicates the enlarged display area. The red line segment represents the data collection area. **B.** Quantification of auxin accumulation shown in panel A. The difference in auxin levels between the *TaPGS1* overexpression lines and wild-type Fielder is significant (n = 10; mean ± SD; * *P* < 0.05; ** *P* < 0.01). **C.** Schematic representation of endosperm cellularization observed through semi-thin sectioning of seeds. The red arrow marks free nuclei in the coenocytic endosperm that has not yet undergone cellularization. **D.** Proportion of seeds with completed cellularization at different DPA. Cellularization was delayed in *TaPGS1* overexpression lines compared to Fielder, particularly at 4.5 to 6 DPA.

To further confirm whether endosperm cellularization is affected, we prepared semi-thin sections to observe seeds at different stages of cellularization. The results showed that in wild-type Fielder, 33.33% of the seeds had completed endosperm cellularization by 4.5 DPA, and all seeds had completed cellularization by 6 DPA. In contrast, none of the *TaPGS1* overexpression seeds had completed cellularization by 4.5 DPA. By 5 DPA, 44.44% of the *TaPGS1* overexpression seeds had completed cellularization, and by 6.5 DPA, all seeds had completed cellularization, indicating a delay of 0.5 days compared to wild-type Fielder (Fig. 4C, D). These results suggest that overexpression of *TaPGS1* causes auxin accumulation in the inner seed coat and delays endosperm cellularization.

### 3.4 Kaempferol treatment recapitulates the cellularization delay

To test whether flavonol accumulation directly affects endosperm cellularization, wild-type seeds at 4 DPA were exposed to increasing concentrations of kaempferol (10, 25, and 50 μM), and cellularization was assessed at 24 and 48 hours. Compared to the control (0 μM), all treated seeds exhibited reduced cellularization, with more pronounced inhibition observed at higher concentrations. Although cellularization progressed further by 48 hours than at 24 hours under each condition, the overall delay was concentration-dependent (Fig. S4). These findings are consistent with the cellularization delay observed in *TaPGS1* overexpression lines, supporting a role for kaempferol in modulating this process.

We next asked whether such a delay affects the final cellular architecture. Seeds from *TaPGS1*-overexpressing and wild-type plants were sampled at 8 DPA, when endosperm cellularization was complete, and prepared semi-thin sections for cell number quantification across three defined endosperm regions (Fig. S3A). *TaPGS1* overexpression lines consistently showed higher cell counts than the wild type. Although region c in line OE-166-6 did not show a statistically significant difference, its cell number was still greater than that of the wild type (Fig. S3B). These results suggest that the delay in cellularization may contribute to an increased endosperm cell number.

### 3.5 *TaPGS1* activates promoters of flavonol pathway genes *TaF3H* and *TaDFR*

To investigate whether the upregulation of *TaF3H* and *TaDFR* in TaPGS1-overexpressing lines results from direct transcriptional activation, we performed transient transactivation assays in tobacco. The promoter regions of *TaF3H* and *TaDFR* were fused to a luciferase reporter (*TaF3H*-pro:LUC and *TaDFR*-pro:LUC), and co-infiltrated with either 35Spro:TaPGS1 or an empty vector control. Luminescence imaging and quantification revealed that TaPGS1 significantly activated both promoters, indicating that TaPGS1 directly promotes the transcription of key genes in the flavonol biosynthetic pathway (Fig. 5A–D).

**Fig. 5.**
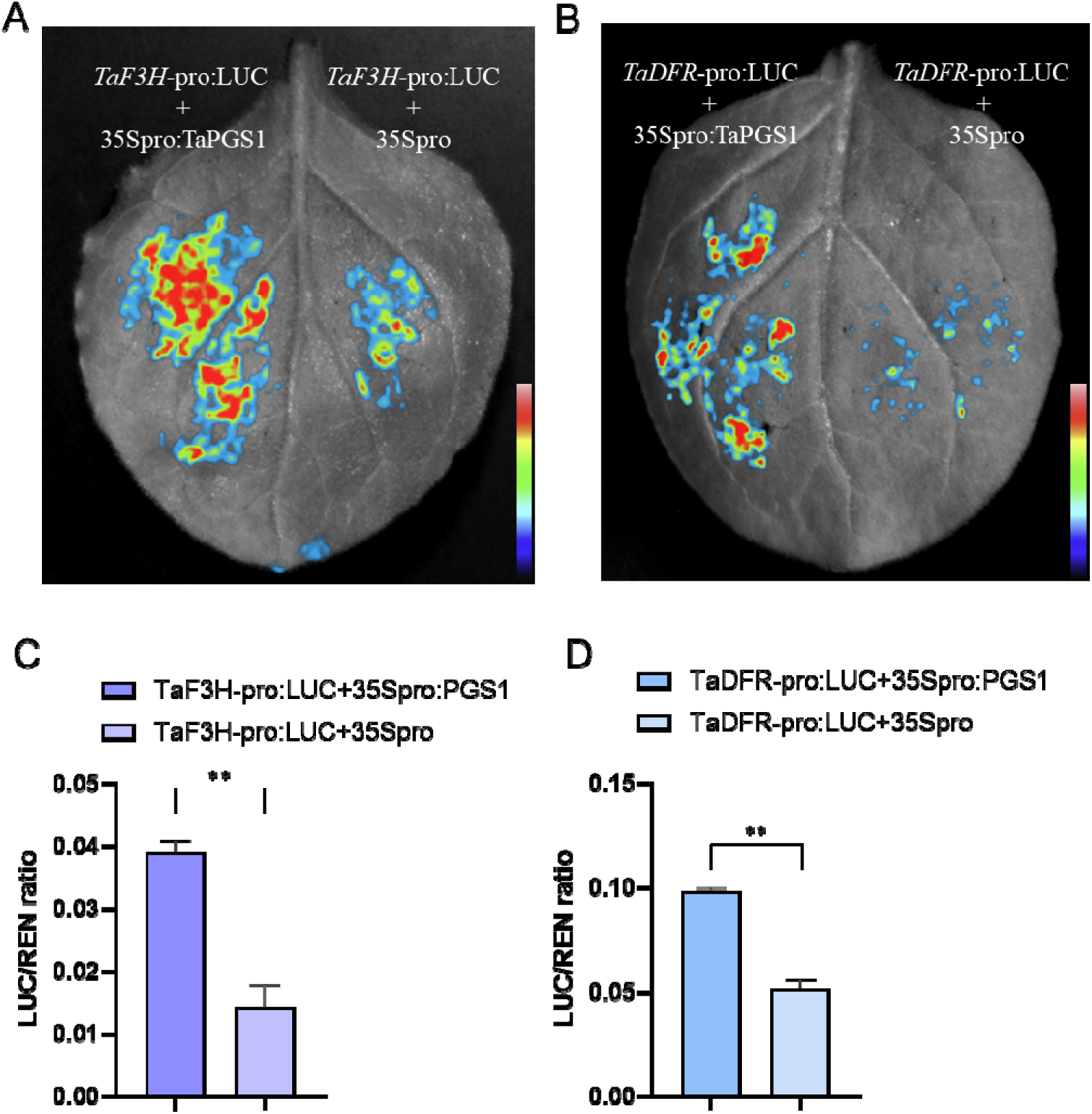
Validation of downstream transcriptional activation of *TaPGS1*. **A.** Validation of transcriptional activation of TaPGS1 with *TaF3H*. The left panel shows the reporter construct *TaF3H*-pro: LUC fused with the 35S promoter and effector *TaPGS1* (35Spro-*TaPGS1*), while the right panel shows the reporter construct *TaF3H*-pro:LUC fused with the empty control (35Spro). B. Validation of transcriptional activation of TaPGS1 with *TaDFR*. The left panel shows the reporter construct *TaDFR*-pro: LUC fused with the 35S promoter and effector *TaPGS1* (35Spro-*TaPGS1*), while the right panel shows the reporter construct *TaDFR*-pro: LUC fused with the empty control (35Spro). C-D. Firefly luciferase (LUC) and Renilla luciferase (REN) activities were measured using the Dual-Luciferase Assay Reagents (Promega)(mean ± SD; * P < 0.05; ** P < 0.01).

Given that bHLH transcription factors are often reported to function in the flavonoid biosynthetic pathway by forming a ternary complex with MYB (Myeloblastosis-related transcription factor) and WD40 (WD40-repeat protein) proteins (Lang et al. 2021; Ramsay and Glover 2005), we investigated whether TaPGS1 might act through similar interactions. Homologs of OsMYB (OsC1) and OsWD40, known to interact with OsRC in rice, were identified in wheat by sequence alignment. Wheat homologs of *OsMYB* were located on chromosomes 4A, 4B, 4D, and 5A, while homologs of *OsWD40* were found on chromosomes 6A and 6B (Fig. S5). Candidate genes with over 90% amino acid similarity were selected for interaction assays. Yeast two-hybrid and BiFC experiments showed that TaPGS1 interacts with TaWD40 but not with TaMYB (Fig. 6A, B), suggesting that TaPGS1 may form a regulatory complex with TaWD40 to activate downstream flavonol biosynthetic genes.

**Fig. 6.**
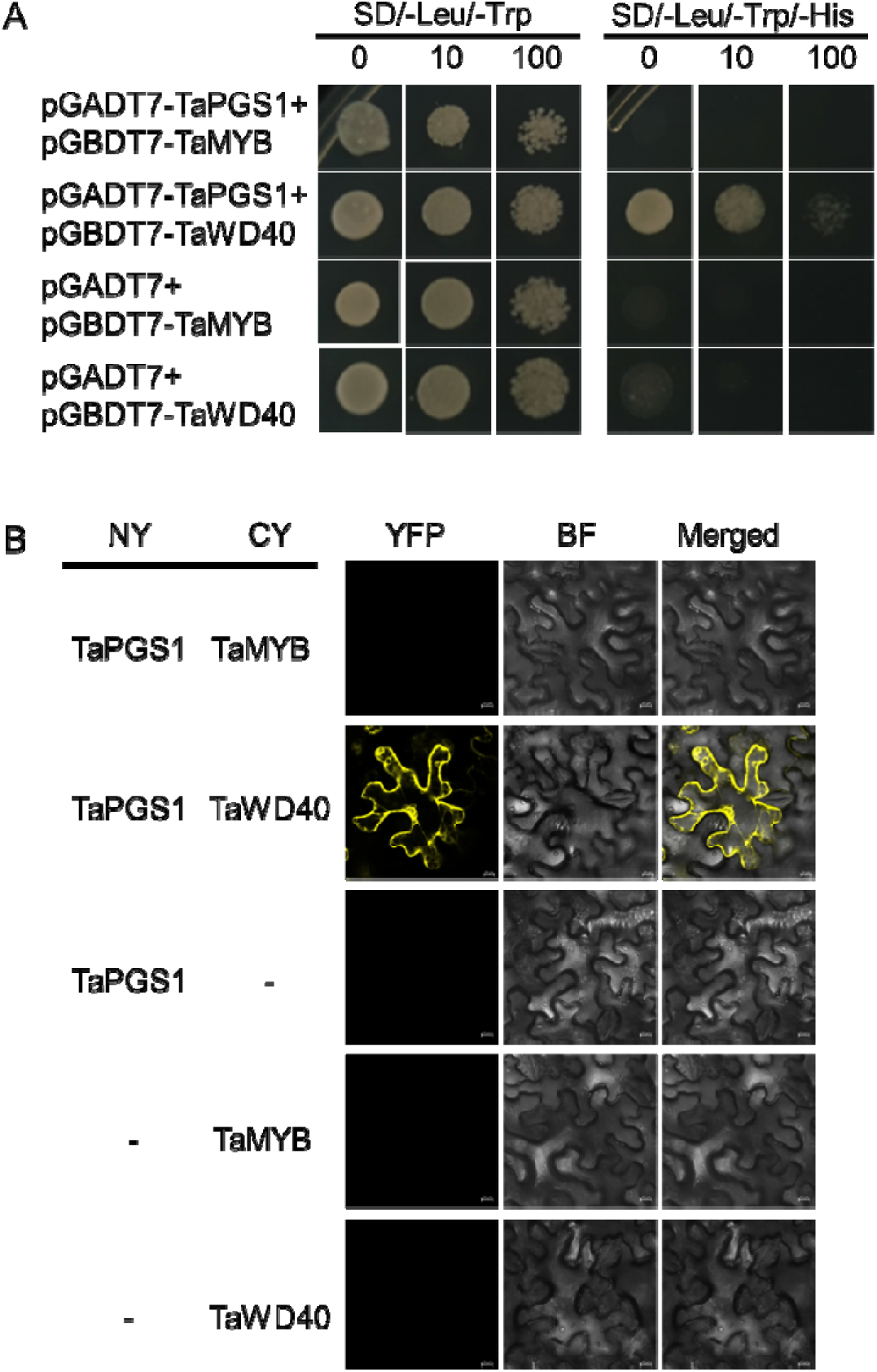
Validation of interaction proteins of TaPGS1. A. Yeast two-hybrid assay validating the interactions between TaPGS1 and TaMYB and TaWD40. B. BIFC validating the interactions between TaPGS1 and TaMYB and TaWD40.

## 4. Discussion

### 4.1 Functional divergence of wheat *TaPGS1* from its *Arabidopsis* and rice homologs

The *Arabidopsis* bHLH protein TT8 and the rice homolog OsRC promote flavonoid and anthocyanin biosynthesis, thereby determining seed-coat pigmentation ( Gu et al. 2011; Nesi et al. 2000). Previous studies have reported that two bHLH homologous genes of *TaPGS1*, *ThMYC4E* (Zhao et al. 2020) and *TaPpb1* (Jiang 2018), are involved in regulating wheat seed color. By contrast, over-expressing *TaPGS1* did not alter the seed-coat color of wheat (Fig. 1A, B), indicating that anthocyanin levels were unchanged.

Transcriptome and metabolite profiling provide a biochemical explanation. Over-expression of *TaPGS1* up-regulated *TaF3H* and *TaDFR* (Fig. 2C; Fig. S2; Fig. S6), increasing the flux from naringenin to dihydrokaempferol. Expression of *F3*′*H* (*Flavonoid 3*′*-Hydroxylase*), both essential for the dihydroquercetin-to-anthocyanin branch, rose slightly but not to a statistically significant level. Although *ANS* transcript abundance rose significantly, the scarcity of the *F3*′ *H*-dependent intermediate eriodictyol meant that insufficient substrate was available for conversion to dihydroquercetin and, ultimately, to anthocyanins (Endo et al. 2012; li et al. 2016; Tian et al. 2015) (Fig. 2C; Fig. S2; Fig. S6). Because flavonol synthase and DFR compete for the same dihydroflavonol substrate, the surplus dihydrokaempferol was preferentially converted to kaempferol, which accumulated in the endosperm, whereas anthocyanin synthesis remained limited (Choudhary and Pucker 2023). Consistent with this preference, the surplus dihydrokaempferol in *TaPGS1* lines was channeled towards kaempferol, which accumulated in the endosperm, while anthocyanin biosynthesis and seed-coat color remained unchanged.

The regulatory context offers a complementary explanation. Anthocyanin production in cereal pericarps requires a complete MYB-bHLH-WD40 (MBW) transcription factor complex. In rice, functional *OsRc* promotes pigmentation only when the structural gene *Rd* (encoding DFR) is active (Gu et al. 2011). In wheat, over-expression of the transcription factor TaMYB10-3D alone is sufficient to darken the normally colorless Fielder pericarp (Lang et al. 2021), demonstrating that Fielder carries functional structural genes for anthocyanin biosynthesis. Our yeast two-hybrid and BiFC assays showed that TaPGS1 interacts with TaWD40 but not with the tested TaMYB protein (Fig. 5D). These findings suggest that, in the Fielder pericarp, TaPGS1 lacks a compatible MYB partner to form the pigmentation-related MYB-bHLH-WD40 complex, whereas other tissues that rely on different combinations of MYB factors may not share this limitation.

Developmental traits further highlight divergence. *OsRc* influences seed dormancy and grain weight in rice, whereas *TaPGS1* over expression enlarged wheat grains without affecting dormancy (Fig. 1C–F). Together, these data show that, despite high sequence similarity, *TaPGS1* has acquired a distinct role in wheat: it redirects flavonoid flux toward flavonol production rather than anthocyanin synthesis and thereby enhances grain size without altering pericarp pigmentation. This divergence likely reflects differences in MBW complex composition and in the availability of downstream structural genes among wheat, rice, and *Arabidopsis*.

### 4.2 Flavonols enlarge wheat grains by impeding auxin efflux and postponing endosperm cellularization

Flavonoids have long been recognized as endogenous inhibitors of polar auxin transport (Jacobs and Rubery 1988). In particular, flavonols such as kaempferol derivatives attenuate PIN-mediated auxin efflux, causing local auxin accumulation (Besseau et al. 2007; Buer and Muday 2004; Peer et al. 2004; Peer et al. 2004; Teale et al. 2021; Yin et al. 2013). Because endosperm cellularization is triggered only after auxin levels fall below a critical threshold (Batista et al. 2019), any restriction of auxin export is expected to delay wall formation.

Our data fits this model. *TaPGS1* overexpression increased kaempferol content in the inner seed coat, reduced auxin export from the endosperm, and led to a marked auxin build-up within the coenocyte (Fig. 3A, B). This auxin surplus delayed the onset of cellularization by ∼0.5 d (Fig. 4A–D), allowing extra rounds of nuclear division and yielding a larger endosperm cell population. Exogenous kaempferol replicated the delay in a dose-dependent manner, confirming that flavonols, rather than *TaPGS1* per se, are the proximate inhibitors. Consequently, the grains from *TaPGS1* overexpression lines were heavier and wider than those of the wild type (Fig. 1D-F).

Although we did not measure auxin flux directly, several lines of evidence argue that the delay reflects inhibited transport rather than increased IAA synthesis. First, transcript levels of auxin-biosynthetic genes (*TAA1*, *YUC*) were unchanged in TaPGS1 lines (Table S5), and pathway-enrichment analysis of the metabolome revealed no up-regulation of auxin-biosynthesis routes (Fig. 2A). Second, immunolocalization showed auxin trapped at the endosperm– seed-coat interface, a distribution pattern typical of blocked efflux. Third, kaempferol is a documented inhibitor of PIN phosphorylation and cycling (Blakeslee et al. 2007; Endo et al. 2020). Together these observations indicate that flavonol-induced transport inhibition leads to auxin accumulation, which in turn postpones cellularization. Nevertheless, radiolabeled IAA transport assays or visualization of PIN dynamics will be required in future work to confirm this mechanistic sequence.

Collectively, these findings support a model in which *TaPGS1* elevates flavonol levels, restricts auxin export from the endosperm, delays cellularization, and thereby permits additional cell divisions that translate into larger wheat grains.

### 4.3 *TaPGS1* regulates seed development and increases seed size through multiple pathways

Previous work demonstrated that overexpression *TaPGS1* in wheat and rice raises thousand-kernel weight and grain size, in part because TaPGS1 activates *TaFL3 / OsFL3* during early grain filling (5–10 DPA) (Guo, et al. 2022). The maize orthologue *ZmFL3* exerts a comparable influence on endosperm storage deposition (Guo, et al. 2022; Li et al. 2017), supporting the view that a *TaPGS1*–*FL3* pathway enhances assimilate accumulation in developing grains.

The present study identifies a second mechanism centered on flavonol metabolism. TaPGS1 up-regulates *TaF3H* and *TaDFR*, increases kaempferol accumulation in the inner seed coat, restricts auxin efflux from the endosperm, and elevates the internal auxin concentration. Auxin build-up postpones cellularization by about half a day, permits additional nuclear divisions, and raises the final endosperm cell number. Exogenous kaempferol repeats this delay in a concentration-dependent manner, confirming that flavonols directly modulate auxin transport. The observed delay is moderate, remaining below the threshold that arrests grain development (Xu et al., 2023), and agrees with reports that brief extensions of the coenocytic phase can enlarge grains (Ando et al. 2023; Zhang et al. 2020).

*TaPGS1* therefore acts through at least two partially independent routes. One enhances assimilate supply via *FL3* activation, and the other expands sink capacity by slowing cellularization through a flavonol–auxin module. Determining the quantitative contribution of each pathway will require genetic separation or temporal regulation of *TaPGS1* targets, yet the convergence of both routes on grain enlargement establishes *TaPGS1* as a promising lever for yield improvement in wheat.

## 5. Conclusion

This study provides a detailed analysis of how *TaPGS1* overexpression in wheat influences seed size by modulating the flavonol pathway and auxin transport. Our results indicate that overexpression of *TaPGS1* leads to the upregulation of genes involved in flavonol biosynthesis, resulting in higher accumulation of flavonols in the seed coat. This accumulation inhibits the transport of auxin from the endosperm, causing auxin to accumulate within the endosperm and delay endosperm cellularization. This delay contributes to an increase in seed size and thousand grain weight. While our findings align with existing literature on the role of flavonols and auxin in seed development, the exact contribution of each pathway to the observed phenotypic changes remains to be fully elucidated. Future research will focus on disentangling these pathways and exploring their combined effects on seed development, with the goal of improving crop yield and quality through targeted genetic manipulation.

## Acknowledgments

This research was funded by the National Key Research and Development Project (2024YFD1201201), the National Natural Science Foundation of China (U22A20472, 32301837), the Technology Support Project of Chengdu (2023-YF08-00008-SN), the Scientific and Technological Innovation 2030 Major Project (2023ZD04069), the Sichuan Science and Technology Support Project (25NSFTD0066), and the open research fund of SKL-CGEUSC (SKL-ZD202212).

## Conflicts of interest statement

The authors declare that they have no competing interests.

## Author contributions

J.W. designed research; X.G. and X.L. performed RNA-Seq and metabonomic analysis; X.G., W.Z., and X.L. performed plasmid construction, genetic transformation, and another assay. X.G., M.J., and X.L. performed phenotypic analysis and planted the experimental material; X.G., X.L. wrote and revised the paper; J.W., H.D., Z.C., and Q.C. supervised the project.

**Supplemental tables Table S1 Primers list.**

**Table S2 Expression levels of 42 flavonoid metabolites in seeds of fielder and *TaPGS1* overexpression lines OE-166-6 and OE-166-39.**

**Table S3 genome-wide gene expression levels (Transcripts Per Million, TPM values) in wheat for Fielder and *TaPGS1* overexpression materials.**

**Table S4 Wheat homologous genes of the anthocyanin biosynthesis pathway ternary complex(bHLH-MYB-WD40).**

**Table S5 The expression levels (TPM values) of auxin-biosynthetic genes in wheat for Fielder and TaPGS1 overexpression materials.**

---

**Fig. S1.**
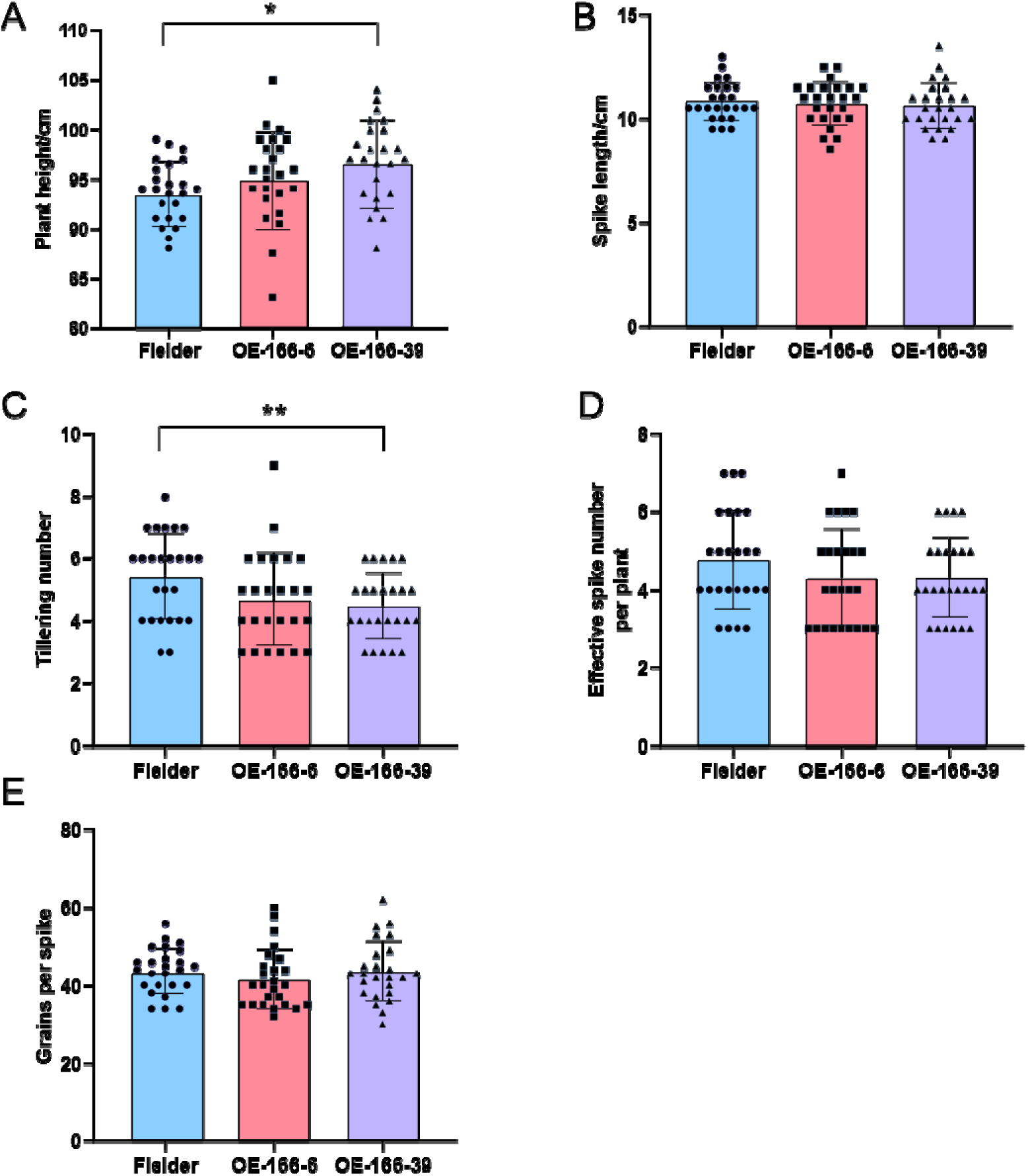
Analysis of key agronomic traits in the field. A-E. Field analysis of Plant height(A), Spike length(B), Tillering number(C), Effective spike(D) and Grains per spike (E) between wild-type Fielder and TaPGS1 overexpression lines. (* *P* < 0.05; ** *P* < 0.01).

**Fig. S2.**
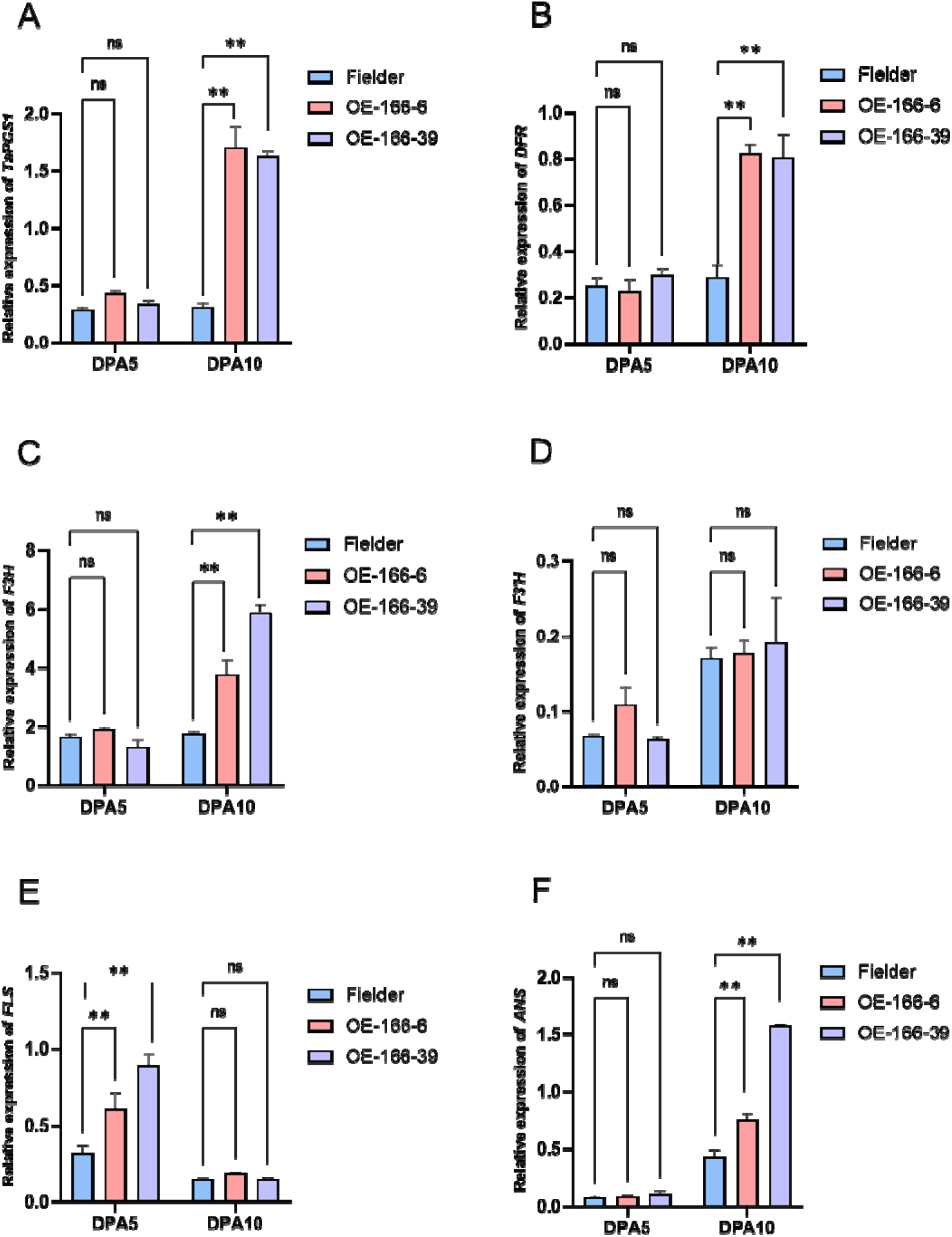
Real-time fluorescence quantitative analysis of TaPGS1 and flavonols pathway-related genes in wild-type Fielder and overexpression lines OE-166-6, OE-166-39. A-F. The relative expression of *TaPGS1* (A), *DFR* (B), *F3H* (C), *F3*’*H*(D), *FLS*(E), *ANS*(F) in wild type Fielder and overexpression lines OE-166-6, OE-166-39 at DPA5 and DPA10 (mean ± SD; * *P* < 0.05; ** *P* < 0.01).

**Fig. S3.**
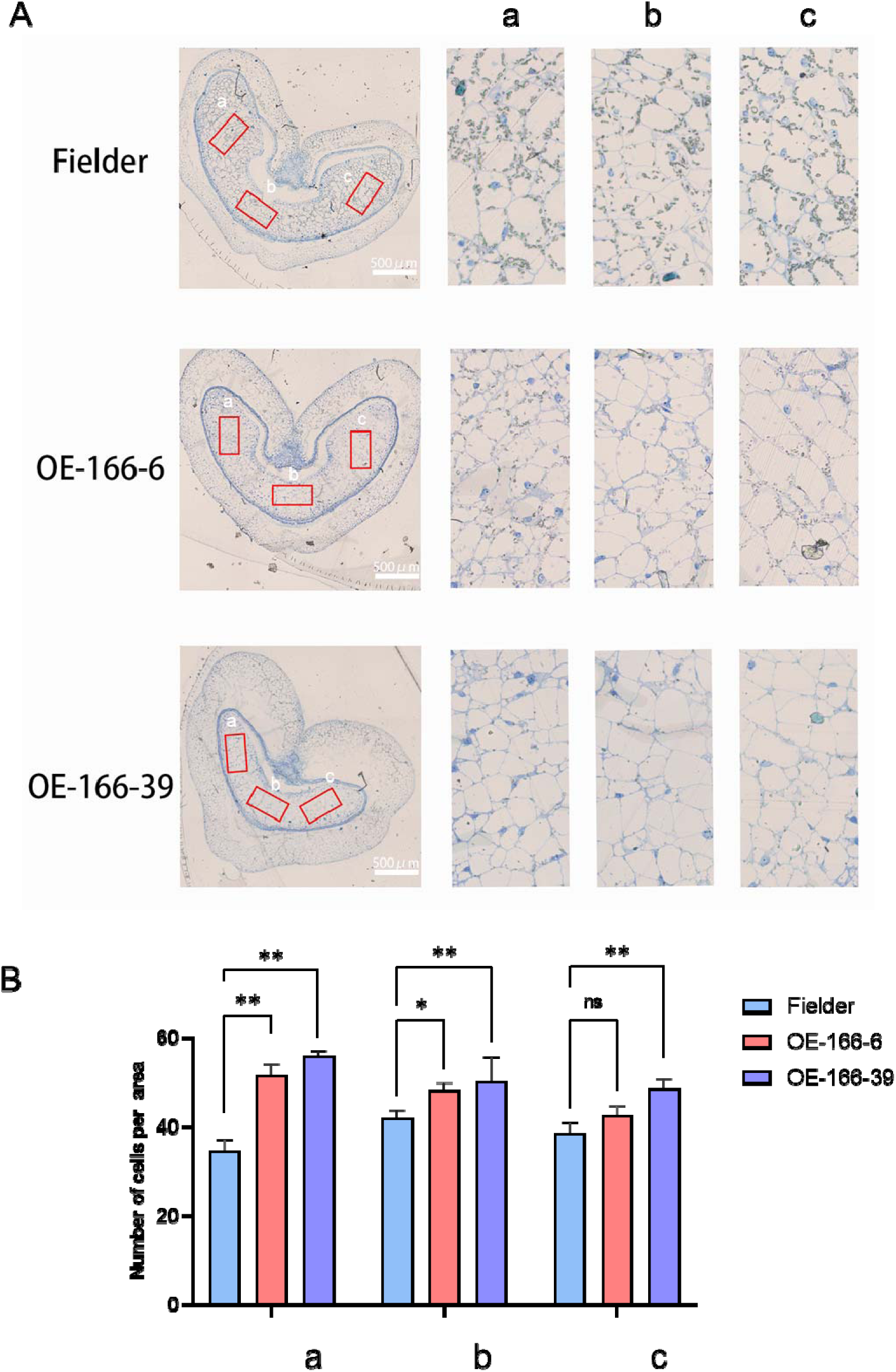
Statistical analysis of endosperm cell number in wild-type Fielder and *TaPGS1* overexpression lines at 8 DPA. **A.** Semi-thin sections showing cell distribution and arrangement in the endosperm of wild-type Fielder and overexpression lines OE-166-6 and OE-166-39 at 8 days post-anthesis (DPA), a stage when cellularization is complete. B. Quantification of endosperm cell number at three defined regions (a, b, and c) indicated in A (n = 10; mean ± SD; * *P* < 0.05; ** *P* < 0.01).

**Fig. S4.**
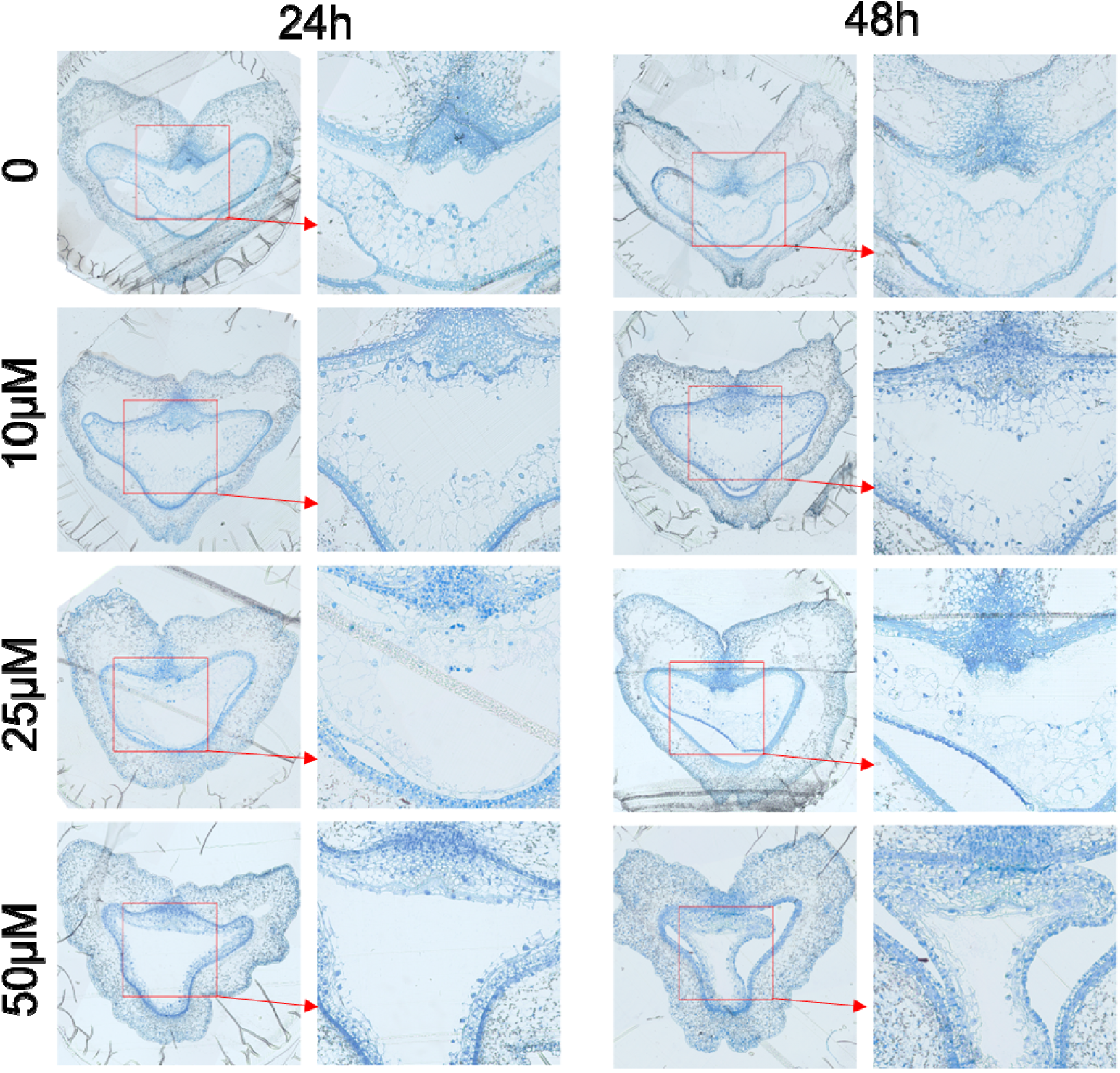
Effects of kaempferol treatment on endosperm cellularization at 4 DPA. Semi-thin sections of wild-type wheat seeds treated with 10, 25, or 50 μM kaempferol for 24 or 48 hours were examined to assess endosperm cellularization. The left panel shows transverse sections after 24-hour treatments; the right panel shows sections after 48-hour treatments. Red boxes indicate regions selected for higher magnification, shown in the corresponding adjacent panels.

**Fig. S5.**
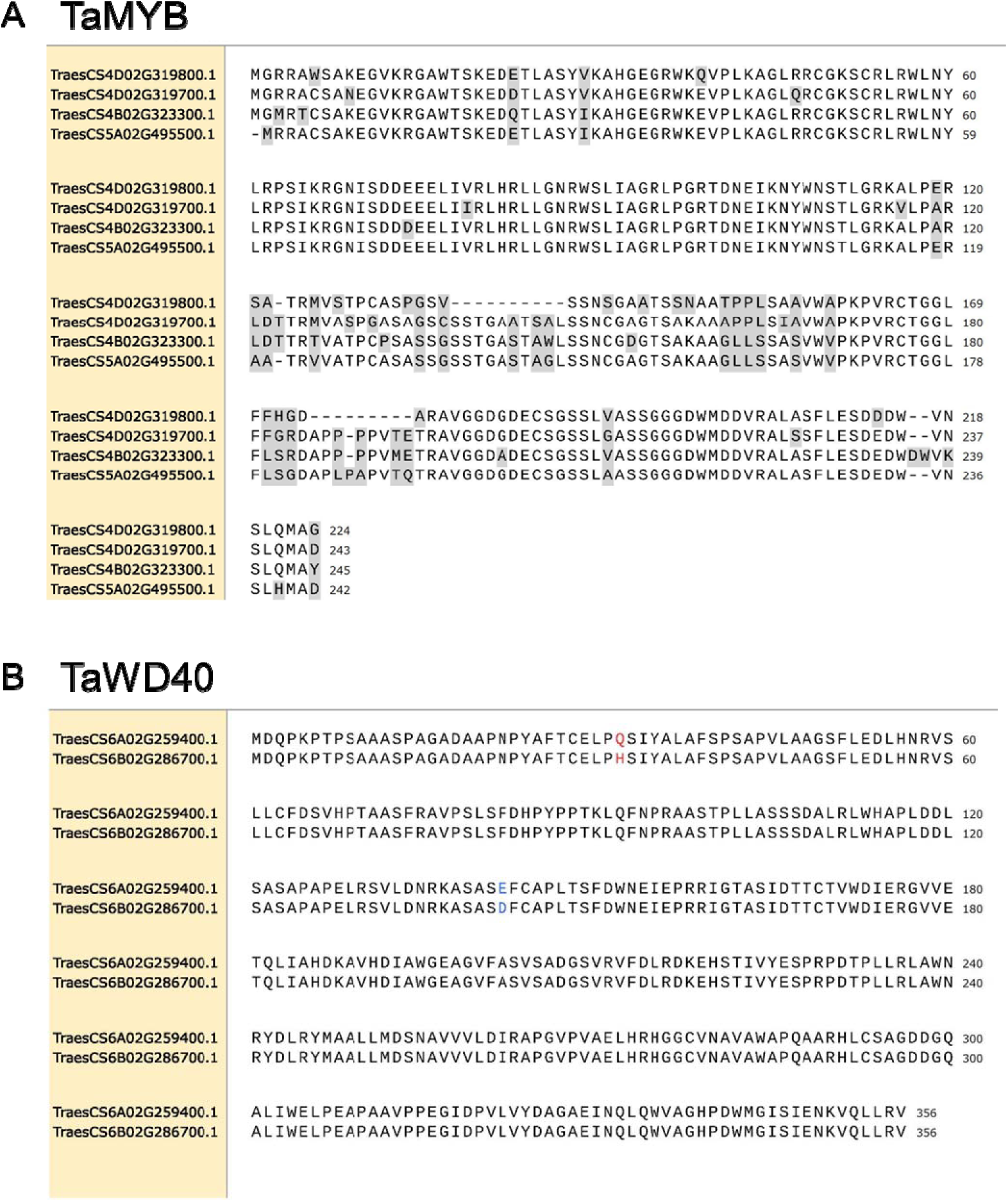
Amino acid sequence alignment of each copy of TaMYB and TaWD40. **A.** amino acid sequence alignment of four copies of TaMYB (*TraesCS4D02G319800.1*, *TraesCS4D02G319700.1*, *TraesCS4B02G323300.1*, *TraesCD5A02G495500.1*). **B.** Amino acid sequence alignment of two copies of TaWD40 (*TraesCS6A02G259400.1*, *Traesc56B02G286700*.1).

**Fig. S6.**
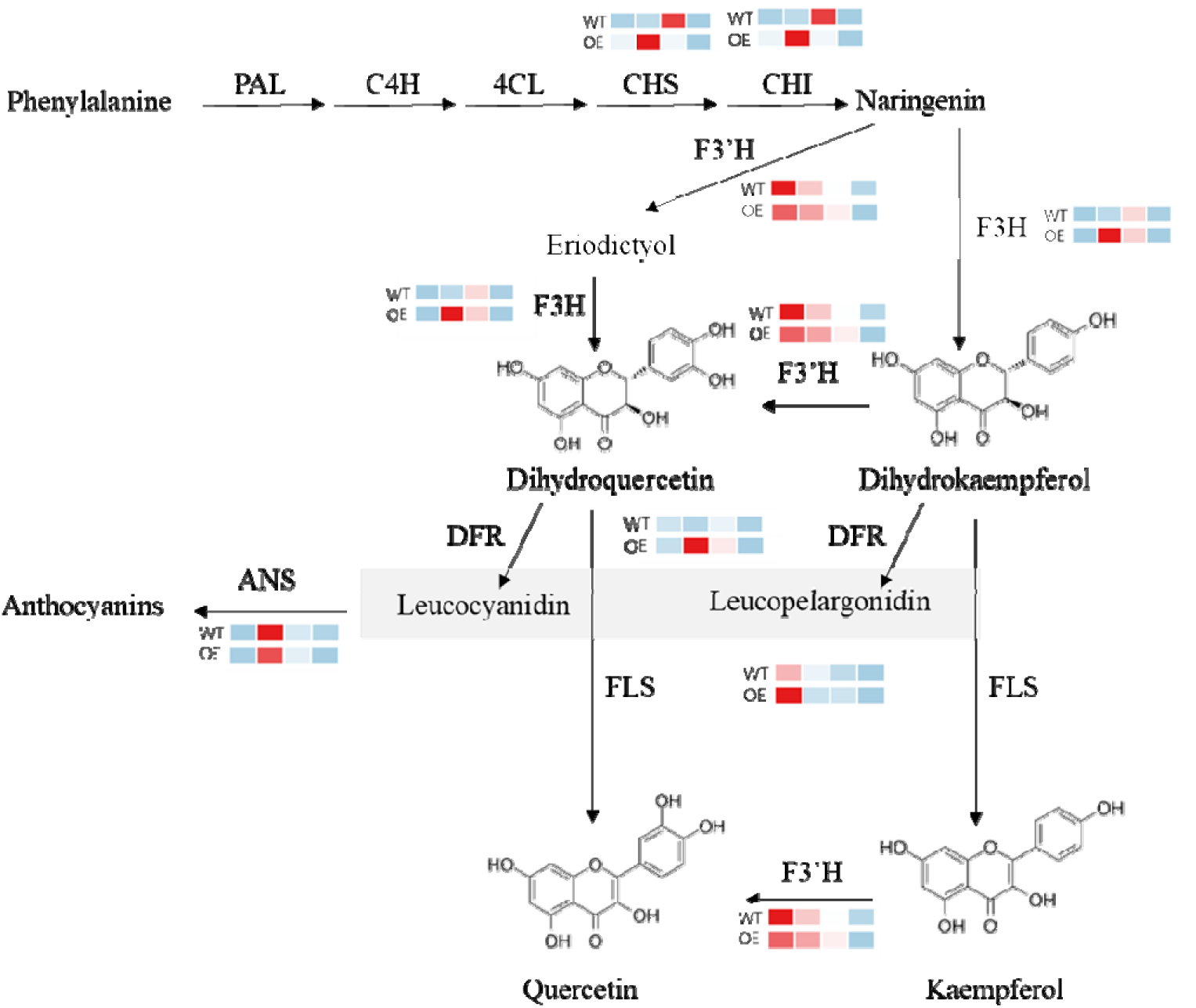
Biosynthetic pathways of flavonols and anthocyanins, and expression profiles of key enzyme genes. The diagram illustrates the metabolic routes of flavonol and anthocyanin biosynthesis, along with transcriptome-based expression levels of key enzyme-encoding genes in *TaPGS1* overexpression lines and wild-type Fielder.

## Notes

### Competing Interest Statement

The authors have declared no competing interest.

